# Deficiency in the autophagy modulator Dram1 exacerbates pyroptotic cell death of Mycobacteria-infected macrophages

**DOI:** 10.1101/599266

**Authors:** Rui Zhang, Monica Varela, Gabriel Forn-Cuni, Vincenzo Torraca, Michiel van der Vaart, Annemarie H. Meijer

**Affiliations:** Institute of Biology Leiden, Leiden University, Einsteinweg 55, 2333 CC, Leiden, The Netherlands

## Abstract

DNA Damage Regulated Autophagy Modulator 1 (DRAM1) is a stress-inducible regulator of autophagy and cell death. DRAM1 has been implicated in cancer, myocardial infarction, and infectious diseases, but the molecular and cellular functions of this transmembrane protein remain poorly understood. Previously, we have proposed DRAM1 as a host resistance factor for tuberculosis (TB) and a potential target for host-directed anti-infective therapies. In this study, we generated a zebrafish *dram1* mutant and investigated its loss-of-function effects during *Mycobacterium marinum* (Mm) infection, a widely used model in TB research. In agreement with previous knockdown analysis, *dram1* mutation increased the susceptibility of zebrafish larvae to Mm infection. RNA sequencing revealed major effects of Dram1 deficiency on metabolic, immune response, and cell death pathways during Mm infection, whereas only minor effects on proteinase and metabolic pathways were found under uninfected conditions. Furthermore, unchallenged *dram1* mutants did not display overt autophagic defects, but autophagic targeting of Mm was reduced in absence of Dram1. The phagocytic ability of macrophages in *dram1* mutants was unaffected, but acidification of Mm-containing vesicles was strongly reduced, indicating that Dram1 is required for phagosome maturation. By *in vivo* imaging we observed that Dram1-deficient macrophages fail to restrict Mm during early stages of infection. The resulting increase in bacterial burden could be reverted by knockdown of inflammatory *caspase a (caspa)* and *gasdermin Eb (gsdmeb)*, demonstrating pyroptosis as the mechanism underlying premature cell death of Mm-infected macrophages in *dram1* mutants. Collectively, these data demonstrate that dissemination of mycobacterial infection in zebrafish larvae is promoted in absence of Dram1 due to reduced maturation of mycobacteria-containing vesicles, failed intracellular containment, and consequent pyroptotic cell death of infected macrophages. These results provide new evidence that Dram1 plays a central role in host resistance to intracellular infection, acting at the crossroad of autophagy and cell death.

## Introduction

Autophagy is an intracellular degradation mechanism that functions to maintain homeostasis and intersects with the initiation of cell death programs when homeostasis is perturbed (1, 2). Autophagy can be induced by various stressors, such as nutrient deprivation and UV damage but also infection. Detection of microbial invaders by the innate immune system activates the autophagy machinery to capture intracellular pathogens in double-membrane autophagosomes and target them to lysosomal degradation (3). Autophagy proteins can also be recruited to single-membrane compartments when pathogens are engulfed by phagocytic cells (3). These autophagic defense mechanisms form promising targets for development of novel host-directed therapies for infectious diseases, many of which are currently complicated by the increasing occurrence of antibiotic resistances (4, 5). This is especially true for tuberculosis (TB), the most lethal infectious disease worldwide. The causative agents of human TB or TB-like disease in poikilothermic animals, *Mycobacterium tuberculosis* (Mtb) and *Mycobacterium marinum* (Mm), are widely studied to increase understanding of the role of autophagy in host defense (3, 6–10).

DNA Damage Regulated Autophagy Modulator 1 (DRAM1) is a stress-inducible regulator of autophagy and cell death. DRAM1 and other members of the DRAM family have been linked to cancer, myocardial infarction, HIV infection and TB, but their molecular and cellular functions remain poorly understood (6, 11–16). Among the five DRAM family members, human DRAM1 was first identified as a p53-induced protein that localizes predominantly to lysosomes and promotes autophagic flux as well as UV-damage induced apoptosis (11). In response to mycobacterial infection, DRAM1 transcription is induced by nuclear factor kappa B (NFκB), a central hub in the signaling network regulating the immune system (6). DRAM1 colocalizes with Mtb in infected human macrophages and is required for host resistance of zebrafish larvae against Mm infection (6). Mtb and Mm share the RD1/ESX1 virulence locus, required to escape from phagosomes into the cytosol (17). Selective autophagy, dependent on ubiquitin receptors such as p62, may counteract this pathogenic mechanism by delaying the escape process or sequestering cytosolic bacteria (3, 8). In addition, selective autophagy has been shown to deliver anti-microbial ubiquitinated peptides to bacteria-containing compartments (18). We have recently shown that selective autophagy receptors are required to control Mm infection in zebrafish and that the host-protective function of Dram1 requires p62 in this infection model (6, 19).

Cytosolic escape of mycobacteria may accelerate the initiation of host cell death programs (8). In TB, the death of an infected macrophage triggers its phagocytosis by other macrophages that subsequently also undergo cell death, resulting in a cascade of cell death events and the formation of inflammatory infection foci, called granulomas (20). The fate of individual infected macrophages is therefore a major determinant of whether granulomas can contain the infection or facilitate dissemination of the infection. It has been previously shown that mycobacteria-infected macrophages can undergo several types of regulated cell death, like apoptosis, necroptosis, and pyroptosis, resulting in different infection outcomes (21). Apoptosis of infected cells is generally regarded as a host-protective defence mechanism against mycobacterial infection, and virulent Mtb therefore actively inhibit apoptosis (20, 22, 23). In contrast, necroptosis and pyroptosis are lytic forms of cell death that create an inflammatory environment that may facilitate extracellular growth and disease progression (24, 25).

While our previous work demonstrated a role for Dram1 in autophagic defense against mycobacterial infection, its potential implication in the regulation of cell death during TB pathogenesis has not been explored. In this study, we generated a *dram1* mutant zebrafish line to address the question. Analysis of the mutant fish showed that Dram1 is required for maturation of Mm-containing vesicles and for macrophages to restrict Mm infection. Without functional Dram1, Mm-infected macrophages prematurelly die via a mechanism dependent on inflammatory Caspase a (Caspa) and Gasdermin eb (Gsmdeb) activities, indicative of pyroptosis. Collectively, our data support that Dram1 protects against mycobacterial infection by modulating autophagic targeting and maturation of Mm-containing vesicles. In the absence of Dram1, infected macrophages rapidly become overburdened by the bacteria and initiate pyroptotic cell death, resulting in increased dissemination of the infection.

## Results

### *dram1* null mutants display increased susceptibility to mycobacterial infection

To study the host resistance function of Dram1, we generated a zebrafish mutant line using CRISPR/Cas9 technology (Fig. S1A). The selected *dram1*^Δ19n/Δ19n^ allele (designated *dram1^ibl53^*) contains a 21 nucleotides deletion combined with a 2 nucleotides insertion in the first coding exon (Fig. 1A, Fig. S1B), which results in undetectable levels of Dram1 protein, supporting that this represents a null allele (Fig. 1B). The mutant was outcrossed to trangenic lines with an autophagy reporter Tg(*CMV:GFP-map1lc3b*) (26) or a macrophage marker Tg(*mpeg1:mCherryF*) (27), hereafter referred to as GFP-Lc3 and mpeg1:mCherry. The offspring from incrossed heterozygous fish (*dram1*^+/Δ19n^) strictly followed Mendelian inheritance (Fig. S1C, D), and homozygous mutants were fertile. Body size measurements indicated no apparent difference in development between *dram1*^Δ19n/Δ19n^ and *dram1^+/+^* larvae (Fig. 1C). Furthermore, the terminal deoxynucleotidyl transferase dUTP nick end labeling (TUNEL) assay did not reveal an effect of *dram1* mutation on the basal level of cell death in zebrafish larvae (Fig. 1D and S1E). In the absence of detectable developmental aberrations, we proceeded to investigate the response of *dram1* mutants to Mm infection. Consistent with previous knockdown results (6), *dram1*^Δ19n/Δ19n^ larvae showed significantly increased susceptibility to infection (Fig. 1E, F). Furthermore, Dram1-deficient larvae displayed accumulation of bacteria inside intersegmental blood vessels, indicative of extracellular bacterial growth (Fig. 1E). We detected no differences in bacterial burden between *dram1^+/+^* and unrelated wild types, indicating that the genetic background did not affect its susceptibility to infection (Fig. 1F). Next, we demonstrated that injection of *dram1* mRNA could rescue the infection susceptibility phenotype of *dram1^Δ19n/Δ19n^*, while injection of a control mRNA containing the Δ19n deletion could not (Fig. 1G). Collectively, our analysis of *dram1*^Δ19n/Δ19n^ zebrafish larvae confirms that Dram1 is necessary for host defense during Mm infection.

**Figure 1:**
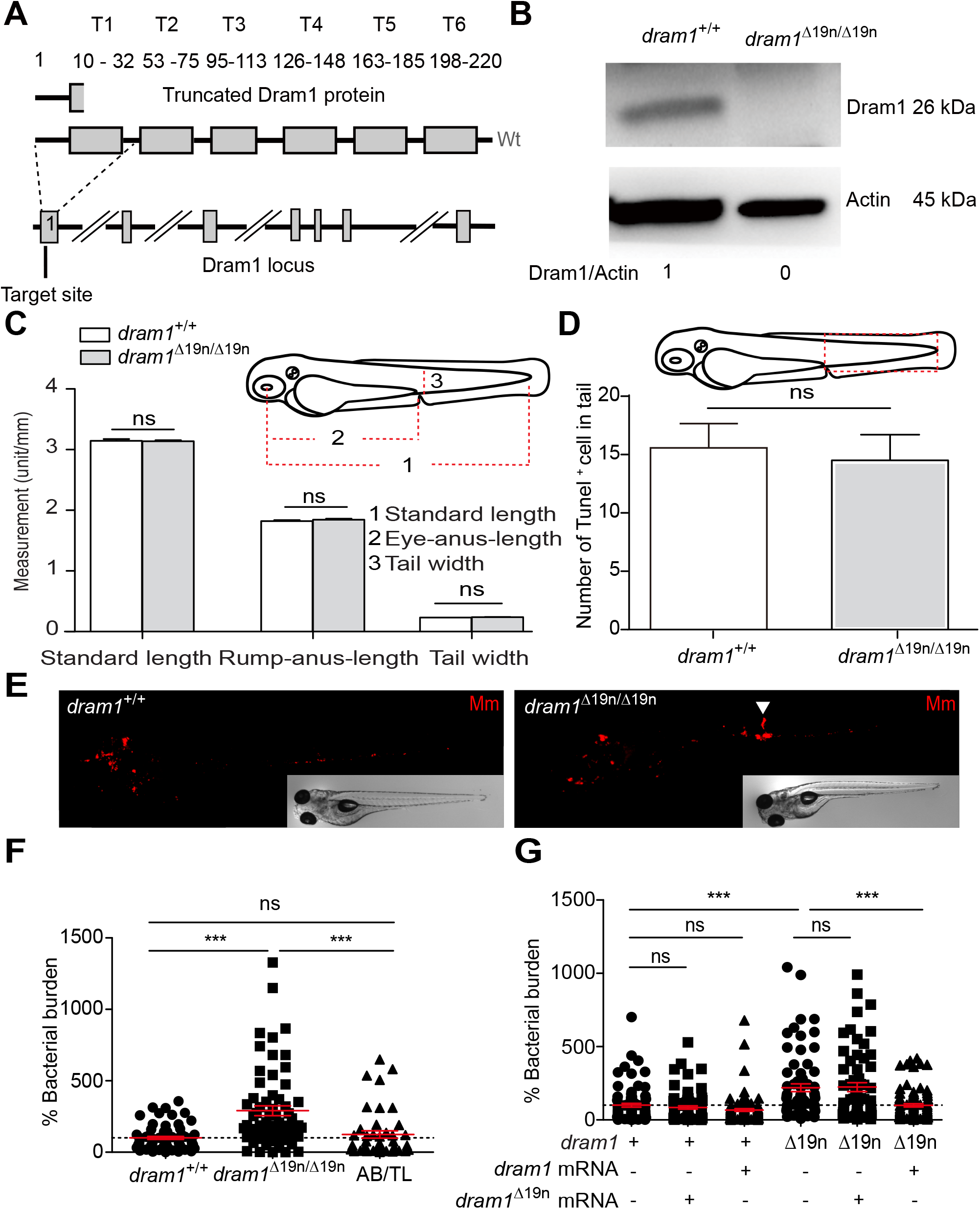
Dram1 deficiency leads to increased susceptibility to Mm infection A. Schematic representation of the zebrafish dram1/Dram1(ENSDARG00000045561/ENSDARP0 0000066996.3) genetic and protein domain architecture and CRISPR/Cas9 target site. Dram1 (240 amino acids) contains six transmembrane domains (indicated with grey boxes and labeled T1-T6 with amino acid numbers indicated above). The gene is depicted with coding exons as grey boxes and introns as solid black lines (introns not drawn to scale). The position of the CRISPR/Cas9 target site and the predicted truncated protein is indicated. B. Confirmation of *dram1* mutation by Western blotting analysis. Protein samples were extracted from 4 dpf *dram1*^Δ19n/Δ19n^ and *dram1*^+/+^ larvae (>10 larvae/sample). The blots were probed with antibodies against Dram1 and Actin as a loading control. C. Measurements of larval body lengths. *dram1*^+/+^ and *dram1*^Δ19n/Δ19n^ larvae (≥10 larvae/group) were imaged by stereo microscopy at 3dpf and body lengths were measured as indicated by the red dotted lines. D. Quantification of TUNEL-positive cells in the indicated region of the tail of *dram1*^Δ19n/Δ19n^ and *dram1*^+/+^ larvae at 3 dpf (≥7 larvae/group). E. Representative stereo images of infected *dram1*^Δ19n/Δ19n^ and *dram1*^+/+^ larvae at 3 dpi. The arrowhead indicates the accumulation of bacteria in intersegmental veins. F and G. Quantification of bacterial burdens at 3dpi for *dram1* mutants, wild type siblings, and unrelated wild types (F) or for *dram1* and *dram1^Δ19n^* mRNA injected individuals (G). The data is accumulated from two independent experiments. Each dot represents an individual larva.

### Dram1 deficiency affects transcriptional regulation of metabolic, immune response, and cell death pathways during mycobacterial infection

To further explore the *dram1*^Δ19n/Δ19n^ phenotype, we performed RNA sequencing. To this end, *dram1*^Δ19n/Δ19n^ larvae were infected with a dosage of 300 CFU, resulting in higher bacterial burden compared with *dram1^+/+^*, or with a lower dosage of 150 CFU, resulting in similar bacterial burden as in *dram1^+/+^* infected with 300 CFU (Fig. 2A, Fig. S2A). The analysis time point, 4 days post infection (dpi), correlates with mycobacterial granuloma formation and strong transcriptional activation of the immune response (28). Principal component analysis showed clear differences between Mm-infected larvae and PBS-injected controls, and between the *dram1*^Δ19n/Δ19n^ and *dram1^+/+^* groups (Fig. S2B). Differential gene expression analysis showed that Dram1 deficiency influences the gene regulation network to a relatively small extent under unchallenged conditions, whereas it has a larger impact on the response to infection (Fig. S2C). Gene ontology and gene set enrichment analysis (GSEA) revealed that *dram1* mutants display an altered transcriptome related to metabolic and proteolytic pathways under non-infected conditions (Table S1). During infection, the transcriptome response of *dram1*^Δ19n/Δ19n^ with 300 CFU showed more overlap with that of *dram1^+/+^* despite the higher bacterial burden. For example, while expression of genes related to host defense pathways, as Nod-like receptor (NLR) signaling, phagosome-related processes, cytokine signaling, and apoptosis, were commonly affected in all Mm-infected larvae, other immune-related pathways, like Toll-like receptor (TLR) and RIG-I-like receptors signaling, were not affected in *dram1*^Δ19n/Δ19n^ larvae infected with similar bacterial burden to *dram1^+/+^* (Fig. 2B). Despite this, approximately 60% of the infection-responsive genes in *dram1^+/+^* (1170 out of 1971) were not differentially expressed in *dram1*^Δ19n/Δ19n^ mutants (Fig. S2D). Alteration of metabolic pathways related to energy and carbon metabolism (e.g. glycolysis, TCA cycle), a characteristic of mycobacterial infections (29), was markedly absent in *dram1*^Δ19n/Δ19n^ larvae (Fig 2B). In contrast, infection of *dram1*^Δ19n/Δ19n^ with 300 CFU influenced expression of genes related to other metabolic processes, as cholesterol and amino acid biosynthesis (Fig. 2B). Further analysis revealed differential expression of several TLRs and downstream genes between the wild type and *dram1* mutant infected groups (Fig. S3). Finally, Dram1 deficiency affected regulation of programmed cell death during infection, resulting in enhanced expression of genes involved in lytic forms of cell death (Fig. 2C).

**Figure 2.**
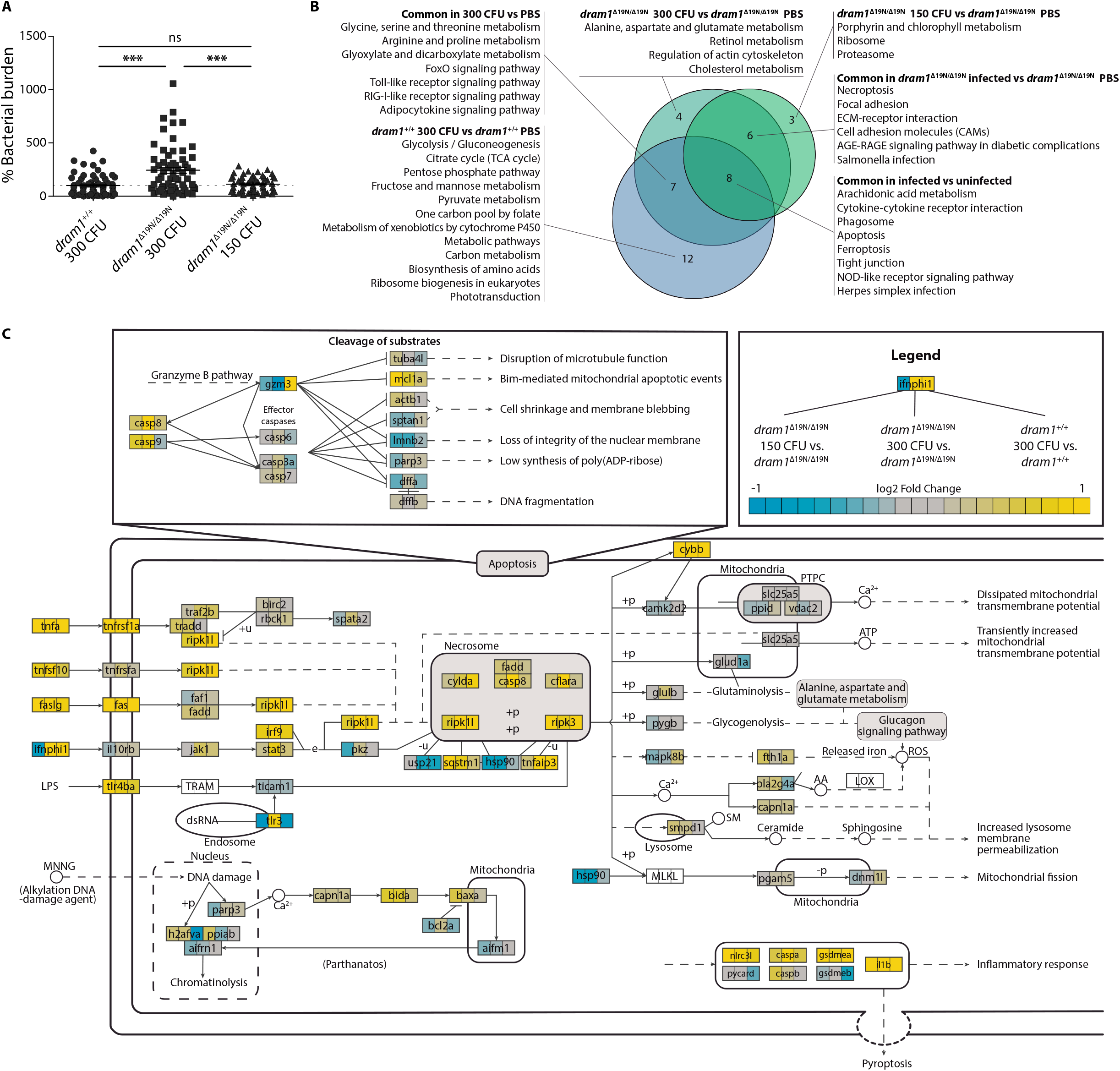
Dram1 deficiency affects gene expression of pathways involved in metabolism, innate immunity and lytic cell death during infection A. Bacterial burdens of larvae injected with different CFU doses of Mm and used for RNAseq. The data is accumulated from three independent sample sets at 4 dpi. Each dot represents the bacterial burden of an individual larva. B. Venn diagram of the significantly enriched KEGG pathways in the transcriptome of larvae infected with Mm. The enrichment comparisons were performed on *dram1*^Δ19n/Δ19n^ 150 CFU versus *dram1*^Δ19n/Δ19n^ PBS, *dram1*^Δ19n/Δ19n^ 300 CFU versus *dram1*^Δ19n/Δ19n^, and *dram1*^+/+^ 300 CFU versus *dram1*^+/+^ PBS. C. Visualization of the lytic cellular death signaling pathway transcriptome shows different responses in the transcriptome of infected *dram1*^Δ19n/Δ19n^ and *dram1*^+/+^. The pathway was adapted from the KEGG pathway necroptosis. In the visualization, the gene expression in the comparison *dram1*^Δ19n/Δ19n^ 150 CFU versus *dram1*^Δ19n/Δ19n^ PBS, *dram1*^Δ19n/Δ19n^ 300 CFU versus *dram1*^Δ19n/Δ19n^, and *dram1*^+/+^ 300 CFU versus *dram1*^+/+^ PBS are depicted by colour gradient (yellow, upregulated, blue downregulated). The expression of all genes of the pathway present in the RNA sequencing was plotted independently of their significance. While the effector genes of the apoptosis pathway did not show high expression changes, the genes from lytic cell death forms, including pyroptosis, showed high expression.

### The autophagic response to Mm infection is altered in *dram1* mutants

Altered metabolic pathway regulation in *dram1* mutants might be a compensatory response to defects in autophagy. Investigating GFP-Lc3 or endogenous Lc3-II accumulation revealed no difference between *dram1*^Δ19n/Δ19n^ and *dram1^+/+^* larvae under unchallenged conditions. However, *dram1*^Δ19n/Δ19n^ larvae responded differently by accumulating higher levels of GFP-Lc3 and Lc3-II compared with *dram1^+/+^* when we applied a cellular stress factor, Bafilomycin A1 (BafA1), which inhibits vacuolar H+ ATPase (V-ATPase) to prevent autophagolysosomal maturation (Fig.3A-C). In agreement, protein levels of ubiquitin-binding receptors, p62 and Optineurin, which are substrates of autophagy (30), were unaffected in unchallenged *dram1*^Δ19n/Δ19n^ larvae, but were elevated compared to the levels in *dram1^+/+^* following BafA1 treatment (Fig. 3D). Similar to BafA1 treatment, Mm infection induced Lc3-II to higher levels in *dram1*^Δ19n/Δ19n^ than in *dram1^+/+^* (Fig. 3E). However, colocalization analysis between GFP-Lc3 and Mm showed that *dram1*^Δ19n/Δ19n^ larvae displayed significantly less GFP-Lc3-positive Mm clusters compared to *dram1^+/+^* (Fig. 3E, F), indicating that autophagic targeting of Mm is reduced in the absence of Dram1, despite an overall increase in Lc3-II accumulation. Taken together, *dram1* mutants display no overt autophagic defects, but are affected in their response to cellular stress, including intracellular infection by Mm.

**Figure 3:**
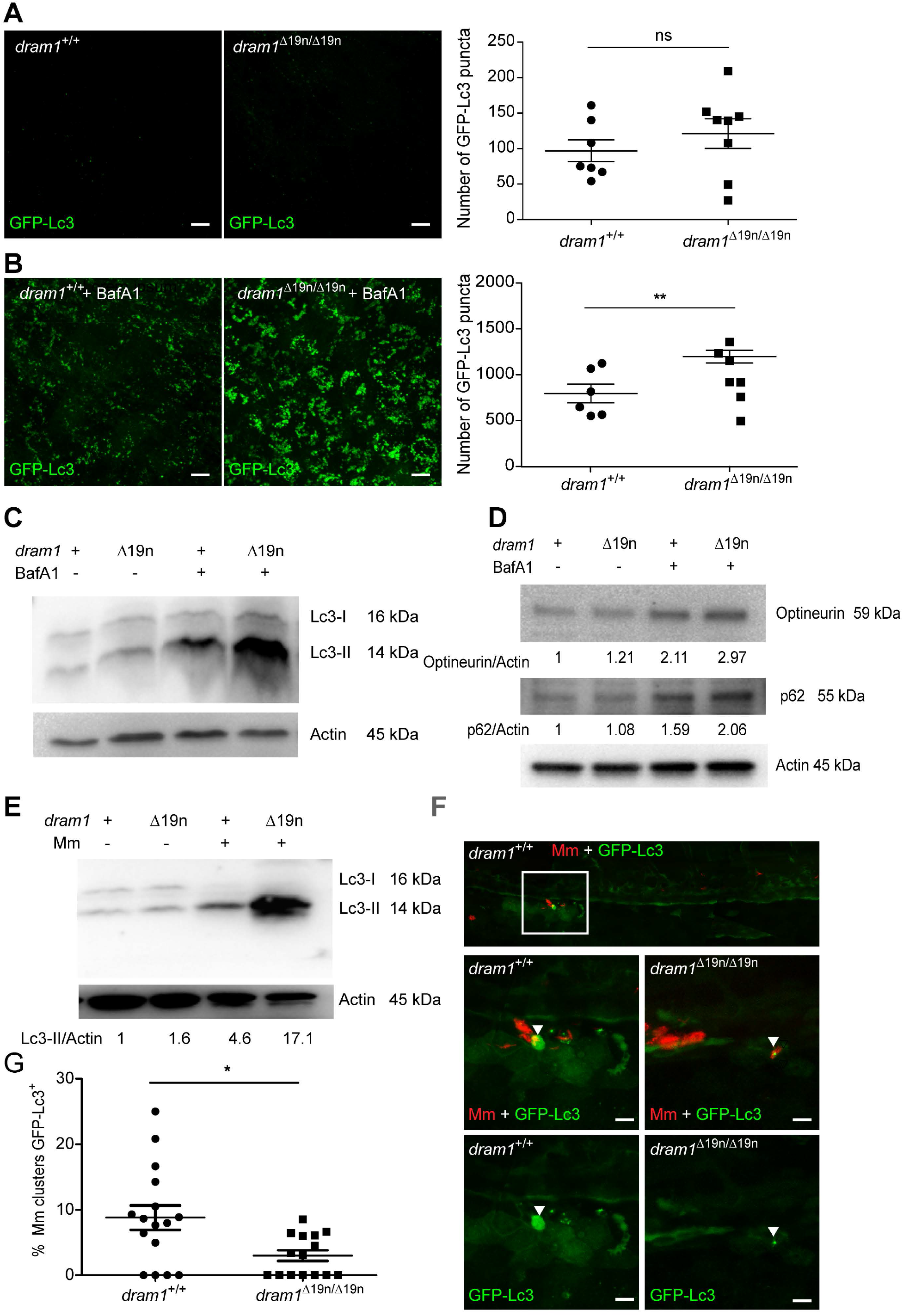
Dram1 is required for GFP-Lc3 targeting to Mm clusters A-B. Representative confocal micrographs and quantification of GFP-Lc3 puncta in *dram1*^Δ19n/Δ19n^ and *dram1*^+/+^ larvae in an unstimulated situation (basal autophagy, A) and following BafA1 treatment (B). Each larva was imaged at a pre-defined region of the tail fin as indicated by the red boxed area in the schematic fish drawing above the graphs. (≥11 larvae/group). Results are accumulated from two independent experiments, each dot represents an individual larva. ns, non-significant *p<0.05,**P<0.01,***p<0.001. Scale bars, 10 μm. C-E. Western blot analysis of autophagy. Protein samples were obtained from 4 dpf *dram1*^Δ19n/Δ19n^ and *dram1*^+/+^ larvae (>10 larvae/sample). Lc3 (C and E), or p62 and Optineurin (D) protein levels were detected in in absence or presence of BafA1 (C and D) or in the presence or absence of Mm (E). Actin was used as a loading control. WB were repeated three (C and D) or two (E) times with protein extracts derived from independent experiments. The Lc3II/Actin or p62/Actin and Optineurin/Actin ratio is indicated below the blots. F-G. Representative confocal micrographs and quantification of GFP-Lc3 co-localization with Mm clusters in infected *dram1*^Δ19n/Δ19n^ and *dram1*^+/+^ larvae. The top image shows the entire region of imaging as indicated in the schematic. The bottom images show GFP-Lc3 colocalization of Mm clusters in *dram1*^Δ19n/Δ19n^ and *dram1*^+/+^ larvae. The arrowheads indicate GFP-Lc3-positive Mm clusters. The data is accumulated from two independent experiments, each dot represents an individual larva (≥15 larvae/group). Scale bars, 10 μm.

### Dram1 deficiency does not affect phagocytosis of Mm

We wanted to exclude that reduced GFP-Lc3 association with Mm might be a consequence of a defect in phagocytosis by macrophages, the primary niche for intracellular Mm growth (31). First, we verified that Dram1 deficiency did not alter the total number of macrophages, labelled by *mpeg1:mCherryF* (Fig. 4A). Similarly, there was no effect on the other main innate immune cell population of zebrafish larvae, the neutrophils (Fig. 4B). Next, we assessed phagocytic activity at 1 h after intravenous delivery of Mm. The results showed that Mm were phagocytosed by macrophages in *dram1^Δ19n/Δ19n^* and *dram1^+/+^* at a similar rate, as indicated by the percentage of intracellular Mm (Fig. 4C). We then determined at which time point during the infection a difference in bacterial burden between *dram1^Δ19n/Δ19n^* and *dram1^+/+^* was first detectable. We found that Dram1 deficiency significantly increased Mm infection burden at 2 dpi but not yet at 1 dpi (Fig. 4E). In conclusion, both *dram1^Δ19n/Δ19n^* and *dram1^+/+^* can phagocytose the injected dose of Mm within the first hour after infection, and the immunocompromised state of Dram1-deficient larvae first becomes apparent two days later.

**Figure 4:**
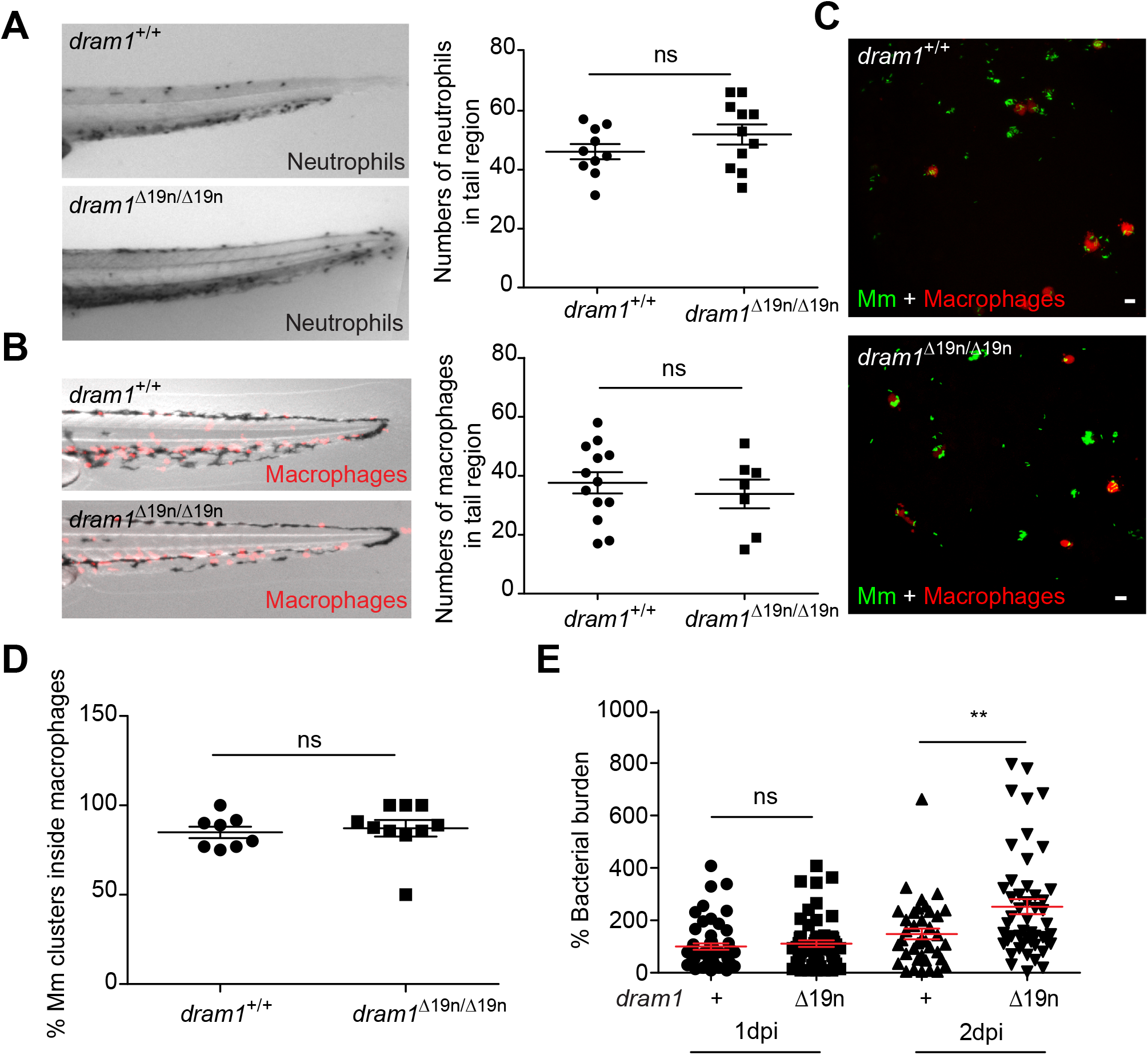
Dram1 deficiency does not affect the capability of macrophages to phagocytose Mm A. Representative stereo images of the whole tail of *dram1*^Δ19n/Δ19n^ and *dram1*^+/+^ larvae following an immunohistochemical peroxidase activity detection protocol. The number of neutrophils in this region was quantified per individual larva (≥18 larvae/group). Each data point represents an individual larva. Results are accumulated from two independent experiments. ns, non-significant *p<0.05,**P<0.01,***p<0.001. B. Representative stereo micrographs of macrophages in the whole tail region and quantification of the number of macrophages in this region. 3 dpf *dram1*^Δ19n/Δ19n^ and *dram1^+/+^/ mpeg1:mCherryF* larvae were obtained from incrossed *dram1*^+/Δ19n^ animals and the number of macrophages for each larva were counted before knowing the genotype. Genotyping was performed by PCR and Sanger sequencing (≥28 larvae/ group). Results are accumulated from two independent experiments, each dot represents an individual larva. ns, non-significant, *p<0.05,**P<0.01,***p<0.001 C. Representative confocal micrographs of the yolk of infected *dram1*^Δ19n/Δ19n^ and *dram1*^+/+^ embryos in *mpeg1:mCherryF* background at 1 hour post infection (hpi). Scale bars, 10 μm. D. Quantification of phagocytosis of Mm by macrophages at 1 hpi. *dram1*^Δ19n/Δ19n^ and *dram1*^+/+^ embryos in *mpeg1:mCherryF* background were infected Mm at 30 hpf and fixed at 1 hpi. Each dot represents the percentage of macrophages that have phagocytosed Mm clusters in an individual larva (≥16 larvae/ group). Results are accumulated from two independent experiments,. ns, nonsignificant, *p<0.05,**P<0.01,***p<0.001. E. Mm bacterial burden for *dram1*^Δ19n/Δ19n^ and *dram1*^+/+^ at 1 and 2 dpi. Each dot represents an individual infected larva.

### Dram1 is required for macrophages to restrict Mm infection

Since Dram1 is a lysosomal membrane protein (11), we asked whether Dram1-deficiency affected the maturation of Mm-containing vesicles. We used LysoTracker to determine the extent of colocalization between Mm and acidic vesicles at an early stage of infection (1 dpi), before differences in bacterial burden were detectable. Approximately 60% of Mm clusters were LysoTracker-positive in *dram1^+/+^*, while this was reduced to 20% in *dram1^Δ19n/Δ19n^* (Fig. 5A, B). Next, we asked whether reduced maturation of Mm-containing vesicles limits the ability of macrophages in *dram1^Δ19n/Δ19n^* hosts to combat the infection. We found that at 1 dpi the majority of Mm clusters were restricted inside macrophages for both *dram1^Δ19n/Δ19n^* and *dram1^+/+^* hosts (Fig. 5C, D). However, at 2 dpi, the majority of Mm (65%) resided inside macrophages in *dram1^+/+^*, while we observed an increased escape of Mm from macrophages in *dram1^Δ19n/Δ19n^* larvae, with only 31% remaining intracellular (Fig. 5E, F). Furthermore, we frequently observed remnants of dead macrophages in the proximity of bacterial clusters in *dram1^Δ19n/Δ19n^* (Fig. 5E). Together, these data demonstrate that Dram1 is necessary for macrophages to contain the infection and prevent extracellular bacterial growth.

**Figure 5:**
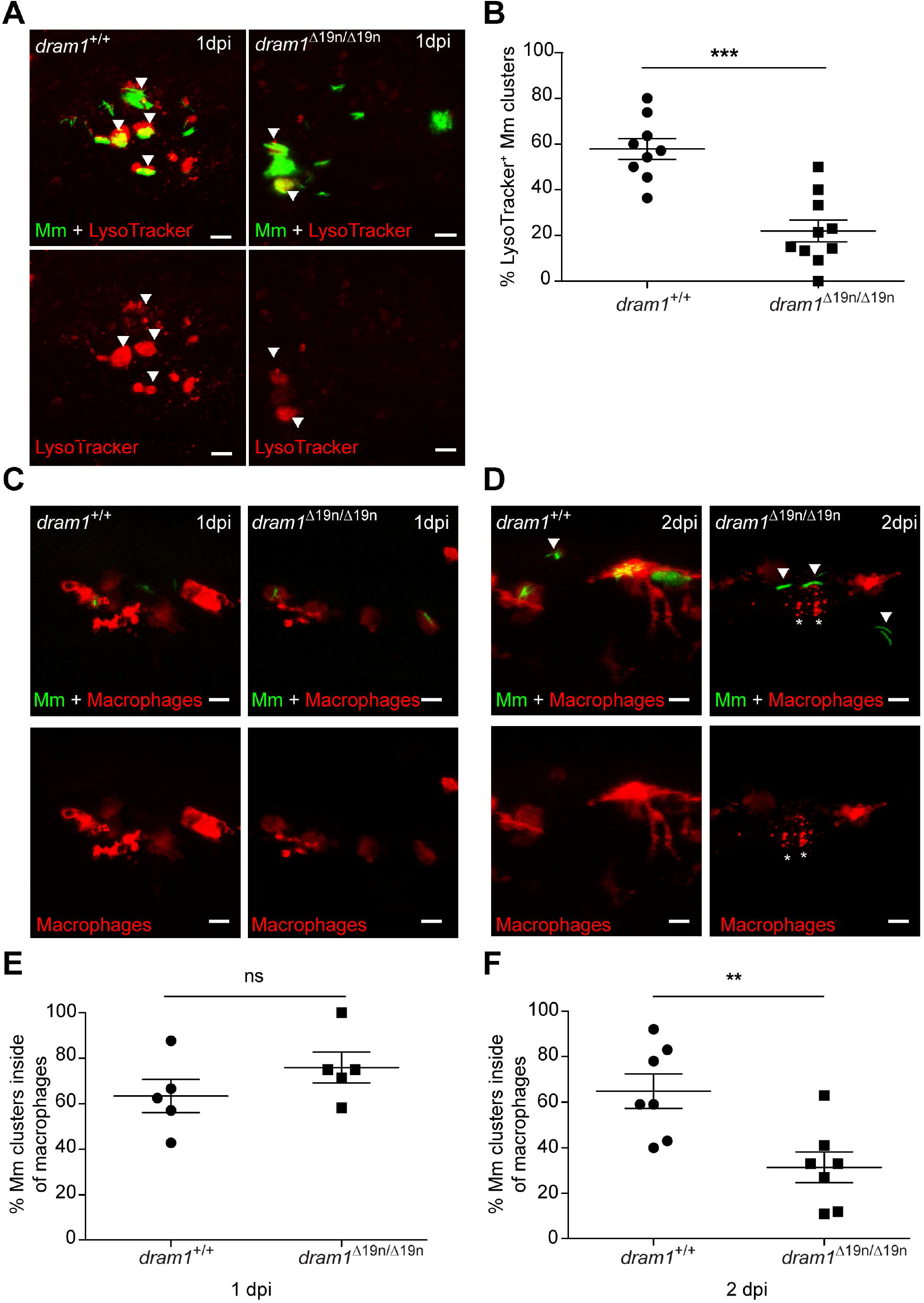
Macrophages fail to restrict Mm infection in Dram1-deficient larvae A. Representative confocal images of LysoTracker staining performed on *dram1*^Δ19n/Δ19n^ and *dram1*^+/+^ embryos at 1 dpi. The arrowheads indicate Lysotracker-positive (LysoTracker^+^) Mm clusters. Scale bars, 10 μm. B. The percentage of LysoTracker^+^ Mm clusters was determined in infected embryos (≥15 embryos/group) at 1 dpi. Each dot represents the percentage of Mm clusters that are LysoTracker^+^ in an individual infected larva. Results are accumulated from two independent experiments. ns, non-significant *p<0.05,**P<0.01,***p<0.001. C and D: Representative confocal images of *dram1*^Δ19n/Δ19n^ and *dram1*^+/+^ embryos/larvae in *mpeg1:mCherryF* background, infected as described in Fig6 A, at 1 dpi (C) and 2 dpi (D). The entire CHT region of fixed embryos or larvae was imaged. The arrowheads indicate extracellular Mm clusters and stars (*) indicate remnants from dead macrophages. Scale bars, 10 μm. E and F: Percentage of Mm clusters restricted inside macrophages at 1 dpi (E) and 2 dpi (F) (≥10 embryos/group). Each dot represents the percentage of intracellular Mm clusters in an individual embryo. Results are accumulated from two independent experiments, ns, non-significant *p<0.05,**P<0.01,***p<0.001

### Dram1 deficiency results in increased pyroptotic cell death of Mm-infected macrophages

Transcriptome analysis had revealed that Dram1 deficiency affects lytic cell death pathways, including necroptosis and pyroptosis, while effects on the apoptosis pathway were relatively minor. To delineate the mechanism responsible for cell death of Mm-infected macrophages in *dram1* mutants, we performed TUNEL staining, which detects damaged DNA present both in apoptotic and pyroptotic cells (32, 33). We observed TUNEL-positive cells around Mm clusters both in *dram1^Δ19n/Δ19n^* and *dram1^+/+^*, but the frequency was around 2.1 times higher in the mutants (Fig. 6A, B). We observed no difference in activation (cleavage) of Caspase 3, a main executioner of apoptosis (34, 35), between *dram1^Δ19n/Δ19n^* and *dram1^+/+^* in the absence or presence of Mm infection (Fig. 6C). Therefore, we asked if the increased cell death of Mm-infected macrophages *dram1^Δ19n/Δ19n^* was due to pyroptosis. Pyroptotic cell death is associated with the activity of inflammatory caspases, like Caspase 1 and Caspase 4/5/11, and is characterized by the formation of Gasdermin pores in the cell membrane (36–38). We have recently found that Caspase a (Caspa) is the Caspase-family member that induces pyroptosis of Mm-infected macrophages in zebrafish via Gasdermin Eb (Gsdmeb) (39). Thus, we analyzed Caspa activity at 2dpf, the time point where we observed increased cell death in *dram1^Δ19n/Δ19n^*. We detected a minor but significant increase of whole larvae Caspa levels in *dram1^Δ19n/Δ19n^* infected with Mm, but not in *dram1^+/+^* (Fig. 6.D). Next, we asked if the increased bacterial burden in Dram1-deficient larvae is dependent on Caspa activity. Knockdown of *caspa* reduced bacterial burden in *dram1^+/+^*. Furthermore, *caspa* knockdown also reduced the enhanced bacterial burden of *dram1^Δ19/Δ19^*, bringing the infection burden in mutants and wild types to a comparable low level (Fig. 6E). Similarly, *gsdmeb* knockdown reduced bacteria burden in *dram1^+/+^* and rescued the hypersusceptibility phenotype of *dram1^Δ19/Δ19^* (Fig. 6F). Collectively, these data suggest that dissemination of mycobacterial infection in zebrafish embryos is promoted in the absence of Dram1 due to the lack of bacterial containment and consequent pyroptosis of infected macrophages.

**Figure 6:**
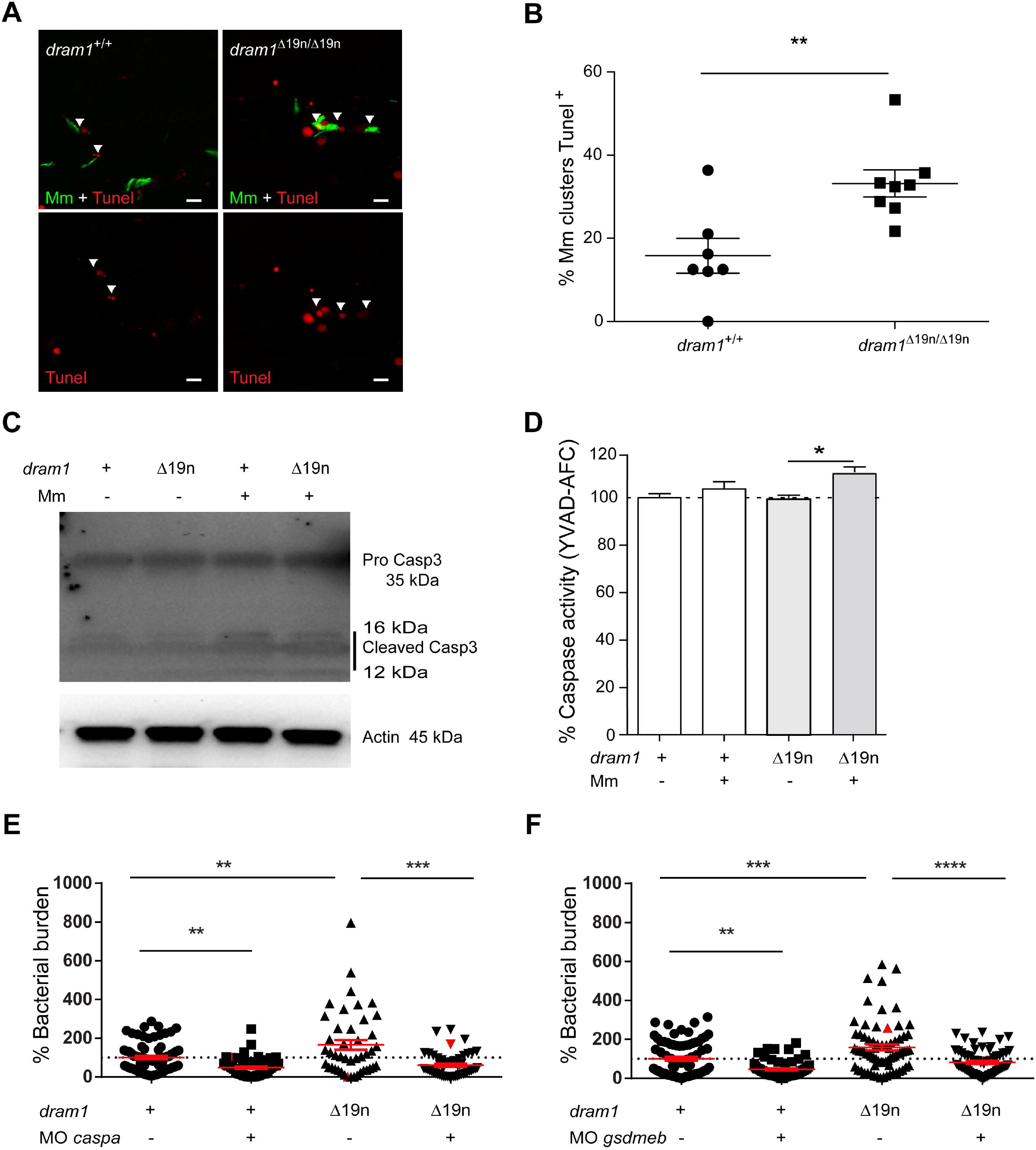
Dram1 deficiency results in increased pyroptotic cell death A. Representative confocal images of TUNEL staining in *dram1*^Δ19n/Δ19n^ and *dram1*^+/+^ larvae at 2 dpi. The entire CHT region of 2 dpi fixed *dram1*^Δ19n/Δ19n^ and *dram1*^+/+^ larvae was imaged. The arrowheads indicate the cells positive for TUNEL staining (TUNEL^+^). Scale bars, 10 μm. B. Quantification of the percentage of Mm clusters TUNEL^+^ in *dram1*^Δ19n/Δ19n^ and *dram1*^+/+^ larvae. Each dot represents the percentage of Mm clusters TUNEL^+^ in the CHT region of an individual infected larva (≥ 12 larvae/group). Results are accumulated from two independent experiments, ns, non-significant,*p<0.05,**P<0.01,***p<0.001 C. Detection of pro-Caspase 3 and cleaved Caspase 3 protein in *dram1*^Δ19n/Δ19n^ and *dram1*^+/+^ embryos. Protein samples were extracted from 4 dpf infected and uninfected *dram1*^Δ19n/Δ19n^ and *dram1*^+/+^ larvae (>10 larvae/sample). The western blots were probed with antibodies against Caspase 3 and Actin as a loading control. Data is representative of two independent experiments. D. Detection of Caspase activity (YVAD-AFC) in *dram1*^Δ19n/Δ19n^ and *dram1*^+/+^embryos. Protein samples were obtained from 2 dpf control and infected *dram1*^Δ19n/Δ19n^ and *dram1*^+/+^ embryos in GFP-Lc3 background (35 embryos/sample). The data is accumulated from two independent experiments, each dot represents an individual larva. ns, non-significant *p<0.05,**P<0.01,***p<0.001 E. Mm bacterial burden at 2 dpi following knockdown of *caspa* in *dram1*^Δ19n/Δ19n^ and *dram1*^+/+^ embryos. The data is accumulated from two independent experiments. F. Mm bacterial burden at 2 dpi following knockdown of *gsmdeb* in *dram1*^Δ19n/Δ19n^ and *dram1*^+/+^ embryos. The data is accumulated from independent experiments, each dot represents an individual larva. ns, non-significant *p<0.05,**P<0.01,***p<0.001

## Discussion

The lysosomal protein DRAM1/Dram1 regulates autophagy and cell survival/death decisions under multiple stress conditions, including diseases like cancer and infection. Its mechanism of action remains largely unknown. Here, we have demonstrated that *dram1* mutation in zebrafish impairs resistance to mycobacterial infection. We show that Dram1 deficiency reduces autophagic targeting of Mm and acidification of Mm-containing vesicles, ultimately resulting in pyroptotic cell death of infected macrophages and increased extracellular growth of mycobacteria during early stages of the infection.

The *dram1* mutant generated for this study was characterized by transcriptome analysis. Under unchallenged conditions, we found that deficiency of Dram1 affects the network of gene regulation to a small degree, with detectable differences in proteinase and metabolic pathways. This transcriptional response could be a compensatory mechanism for defects in lysosomal function due to the deficiency in Dram1. This hypothesis is in line with recent studies that have revealed that lysosomes function as central regulatory units in signal transduction (40). The differences between *dram1* mutants and wild types in the expression of metabolic pathway genes was markedly enhanced in response to Mm infection. Many recent studies have shown that the metabolic status of macrophages is critical for their innate host defense function (41) Therefore, it is conceivable that metabolic dysregulation in *dram1* mutants is a major factor in the hypersusceptibility phenotype. However, this phenotype may also depend on altered regulation of innate immunity signaling pathways that play an essential role in resistance to infection, including TLR signaling. Differential expression of cell surface or endosomal members of the TLR pathway could reflect a response to an increased cytosolic or extracellular localization of Mm in *dram1* mutants. Finally, upregulated expression of genes in lytic cell death pathways may contribute to pathological inflammation and is consistent with the increased bacterial burden in *dram1* mutants.

The function of DRAM1 as a modulator of autophagy has been studied well *in vitro* (11). We therefore tested whether zebrafish *dram1* mutants display defects in autophagic processes. Autophagy is a host response to diverse stress factors, including starvation. Zebrafish larvae until 5 dpf can rely on their yolk proteins for nutrients (42), and we therefore assumed that their autophagic processes are not activated above a level normal for their developmental stage, unless autophagy is triggered by a stressor such as infection. In agreement, we did not detect any differences when comparing the basal levels of autophagy activity in uninfected *dram1* mutant larvae of 4 dpf to those of their wild type siblings. This finding is consistent with an *in vitro* study of the function of mouse DRAM1 (43), which showed that basal autophagy was not altered in the absence of DRAM1 in primary mouse embryonic fibroblasts (MEFs). The five members of the DRAM family are conserved between human, mouse and zebrafish (6, 11, 44–47). Therefore, it is conceivable that other DRAM family members can replace the loss of Dram1/DRAM1 under basal conditions, or that DRAM1 is only involved in autophagic processes in response to specific stress factors. DRAM1 deficiency did not change autophagy induction in response to starvation or while blocking mTOR (48). However, the lack of DRAM1 affected the activation of autophagy in human cells (HeLa and A549) following the induction of cellular stress by treatment with the mitochondria inhibitor 3-nitropropionic acid (3-NP) (49). Besides infection, DNA-damage, and interference with energy metabolism (6, 11, 14), it remains to be further investigated which stress factors can activate DRAM1/Dram1 *in vitro* and *in vivo*.

Overnight treatment of *dram1* mutant larvae with BafA1, which blocks lysosomal degradation of autophagosomes, revealed an increase of GFP-Lc3 puncta and Lc3-II protein levels. The *dram1* mutants also accumulated higher Lc3-II protein levels than their wild type siblings under conditions of Mm infection. It is possible that the prolonged stress conditions imposed to zebrafish larvae during BafA1 treatment or infection induce a compensatory response in *dram1* mutants to produce more autophagosomes. Importantly, despite of the increased Lc3-II levels in infected *dram1* mutants, imaging in GFP-Lc3 transgenic fish revealed that mycobacteria are targeted by autophagic vesicles nearly 3-folds less frequently in *dram1* mutants compared to wild type zebrafish larvae. This reduced autophagic targeting of Mm was not due to a different phagocytic ability of zebrafish macrophages. We did, however, find that Dram1 deficiency reduced acidification of Mm-containing vesicles, which was associated with premature death of infected macrophages. The resulting higher mycobacterial burden of infected zebrafish is in line with increased expansion of extracellularly growing Mm (50). While we found that Mm-infected macrophages die more rapidly in the absence of Dram1, human DRAM1 has been shown to trigger lysosomal membrane permeabilization and cell death in HIV-infected CD4+ T cells, thereby lowering viral replication (14). Together, these studies indicate that DRAM1/Dram1 expression levels can have a major impact on cell death processes during infection, with different outcomes dependent on the cell type and infectious agent.

DRAM1 was previously shown to mediate apoptosis by blocking the degradation of the pro-apoptotic protein Bax (12). While Dram1 deficiency leads to more cell death during Mm infection of zebrafish larvae, we did not observe any changes in total cell death based on TUNEL assays and CASP3 processing. In addition, transcriptome analysis revealed more pronounced effects on genes involved in lytic types of regulated cell death. Strikingly, we found that Dram1 deficiency leads to increased inflammatory caspase activity and gasdermin-dependent pyroptotic cell death. Previous studies revealed that pyroptosis can be induced by diverse pathogens and forms a critical mechanism to restrict microbial infection (51, 52). In line with this, there is also evidence that mycobacteria inhibit pyroptosis of infected macrophages via diverse mechanisms (53). However, recent studies found that lytic cell death (pyroptosis and necrosis/necroptosis) helps mycobacteria to evade host immunity and disseminate the infection (25, 54). Indeed, in the present study we found that pyroptotic cell death promotes the expansion of mycobacteria in Dram1-deficient zebrafish hosts. Moreover, genetic inhibition of this cell death pathway could rescue the exacerbated bacterial growth in *dram1* mutants. Taken together, the death of infected macrophages is intricately related to TB pathogenesis and can result either in increased dissemination or restriction of the infection in the zebrafish host. The contradicting evidence discussed above concerning the beneficial or detrimental effects of the different modes of cell death suggests that the balance between different cell death modalities plays a crucial role in determining the outcome of the infection and a carefull characterization of the specific type of cell death being studied is critical for a better understanding of TB pathogenesis.

In conclusion, restriction of mycobacteria in infected macrophages during the early stages of infection requires functional Dram1. In this work, we have shown that Dram1 is involved in several processes important to defense against intracellular pathogens, potentially providing an intersection between modulation of autophagy, lysosomal function, and programmed cell death. Future studies are required to precisely elucidate the role of the lysosomal protein Dram1/DRAM1 in this network. Facing the rise in multidrug resistant Mtb strains, there is an urgent need to improve treatment strategies to control TB progression. Host-directed therapies have emerged as a promising alternative to counter TB. Drugs targeting host defense mechanisms or processes such as vesicle trafficking can assist the host in responding appropriately to Mtb infection, thereby promoting the effectiveness of drug treatments and reducing the time required for treatment (4, 5). Using an *in vivo* model for the early stages of TB disease we have demonstrated the importance of Dram1 for the elimination of intracellular mycobacteria and the cell fate of infected macrophages. This makes DRAM1/Dram1 – and its interaction partners that remain to be identified – promising drug targets to improve the outcome of TB disease.

## Materials and methods

### Zebrafish culture and lines

Zebrafish lines in this study (Table S2) were handled in compliance with local animal welfare regulations as overseen by the Animal Welfare Body of Leiden University (License number: 10612) and maintained according to standard protocols (zfin.org). Generation of the *dram1* mutant was approved by the Animal Experimention Committee of Leiden University (UDEC) under protocol 14198. All experiments were done on embryos or larvae up to 5 days post fertilization, which have not yet reached the free-feeding stage. Embryos/larvae were kept in egg water (60 ug/ml Instant Ocean sea salts) containing 0.003% 1-phenyl-2-thiourea (PTU, SIGMA-ALDRICH) at 28.5°C and treated with 0.02% ethyl 3-aminobenzoate methanesulfonate (Tricaine, SIGMA-ALDRICH) for anesthesia before bacterial injections, imaging and fixation.

### CRISPR/Cas9 mediated mutagenesis of zebrafish *dram1*

Short guide RNAs (sgRNAs) targeting the first coding exon of zebrafish *dram1* (ENSDARG00000045561) were designed using the chop-chop website (55) and generated by PCR complementation and amplification of full length ssDNA oligonucleotides (Sigma-Aldrich, Table S3) as described (19). For *in vitro* transcription of sgRNAs, 0.2 μg template DNA was used to generate sgRNAs using the MEGA short script ^®^T7 kit (AM1354, ThermoFisher) and purified by RNeasy Mini Elute Clean up kit (74204, QIAGEN Benelux B.V., Venlo, Netherlands). The Cas9 mRNA was transcribed using mMACHINE^®^ SP6 Transcription Kit (AM1340, ThermoFisher) from a Cas9 plasmid (39312, Addgene) (Hrucha et al 2013) and purified with RNeasy Mini Elute Clean up kit (74204,QIAGEN Benelux B.V., Venlo, Netherlands). A mixture of sgRNA and Cas9 mRNA was injected into one cell stage AB/TL embryos (sgRNA 150 pg/embryo and Cas9 mRNA 300 pg/embryo). The effect of CRISPR injection was confirmed by PCR and Sanger sequencing. Genotyping was performed by PCR-amplification of the genomic region of interest using the following primers: Forward: 5’-AGTGAACGTCCGTGTCTTTCTT-3’, Reverse: 5’-ACATCTTGTCGATACAAAGCGA-3’; followed by Sanger sequencing to identify mutations (Base Clear, Netherlands) (19).

### RNA sequencing

Total RNA was extracted from 5 dpf infected and non infected snap-frozen larvae (20 larvae/sample) from three independent crosses using Qiazol reagent (79306, QIAGEN) according to the manufacturer’s instructions and extracted with RNeasy Mini kit (74104, QIAGEN). RNAs were quantified using a 2100 Bioanalyzer (Agilent, US). At least 10 million reads per sample were sequenced using Illumina Single read 50 nt runs in a Hiseq2500. Sequencing, mapping the reads against the *D. rerio* GRCz10.80 reference genome and read counting were performed by ZF-screens (Leiden, Netherlands). Analysis of the count libraries was performed in RStudio 1.1.383 (56) running R 3.4.3 (57) using in-house scripts (available at github.com/gabrifc). An initial quality check of the samples was performed using the tools provided in the edgeR package v3.20.7 (58). Differential gene expression was assessed via pairwise comparisons using DESeq2 v1.18.1 (59). Genes with a FDR-adjusted p-value (adjpval) < 0.05 were considered statistically significant. Venn Diagrams were created using the R package VennDiagram v1.6.18 (60). Gene lists were ranked using the published function “-log_10_(adjpval)*log_2_(fold-change)”, compared to the C2 “Curated Gene Sets” collection from the Molecular Signatures Database (MSigDB) using GSEA v3.0 (61), and visualized with fgsea v1.4.1 (62). Gene ontology enrichment was analysed with goseq v1.3.0 (63). Updated gene length and Gene Ontology data from the Zv10 assembly was retrieved from Ensembl with the packages ensembldb v2.2.1 (64) and biomaRt v2.34.2 (65), respectively. When necessary, mapping between different database gene identifiers was also performed using biomaRt v2.34.2. KEGG pathway analysis was performed with the kegga function provided in limma v3.34.5 (66). Gene regulation data of significant pathways was visualized with pathview v1.18.0 (67). Data are deposited into Gene Expression Omnibus under accession number GSE129035.

### Western blot analysis

Western blot analysis was performed as previously described (19) Antibodies used were as follows: polyclonal rabbit anti DRAM1 (N-terminal) (1:1000,ARP47432-P050, Aviva systems biology), polyclonal rabbit anti-Optineurin (C-terminal) (1:200, lot#100000; Cayman Chemical), polyclonal rabbit anti-p62 (C-terminal) (1:1000, PM045, lot#019, MBL), polyclonal rabbit anti Lc3 (1:1000, NB100-2331, lot#AB-3, Novus Biologicals), monoclonal Caspase 3 antibody (1:1000, #9662, Lot#12, Cell Signaling), Anti mono-and polyubiquitinated conjugates mouse monoclonal antibody (1:200; BML-PW8810-0100, lot#01031445, Enzo life Sciences), Polyclonal actin antibody (1:1000, 4968S, lot#3, Cell Signaling), Anti-rabbit IgG, HRP-Linked Antibody (1:1000, 7074S, Lot#0026, Cell Signaling), Anti-mouse IgG, HRP-linked Antibody (1:3000, 7076S, Lot#029, Cell Signaling).

### Infection conditions and bacterial burden quantification

*Mycobacterium marinum* strain M or *Mycobacterium marinum* strain 20 fluorescently labeled with mWasabi or mCherry, respectively (68, 69), were microinjected into the blood island of embryos at 28 hpf as previously described (70). The injection dose was 200 CFU for all experiments, except for the phagocytosis assay (500 CFU), and the RNA sequencing of infected wild types (300 CFU) and *dram1* mutants (150 and 300 CFU). Embryos were manually dechorionated by tweezers before the injection. Infected embryos were imaged using a Leica MZ16FA stereo fluorescence microscope equipped with a DFC420C color camera, and the bacterial pixels per infected fish data were obtained from the individual embryo stereo fluorescence images using previously described software (71).

### Confocal laser scanning microscopy and image quantification

Fixed or live embryos were mounted with 1.5% low melting agarose (140727, SERVA) and imaged using a Leica TCS SPE confocal microscopy. For quantification of numbers of GFP-Lc3 positive vesicles, the fixed 4dpf larvae were imaged by confocal microscopy with a 63x water immersion objective (NA 1.2) in the pre-defined tail fin region to detect the number of GFP-LC3-positive vesicles (Fig3 B and C). The numbers of GFP-Lc3 vesicles were measured by Fiji/ImageJ software (Fig. 3 B, C) (72). For quantification of the autophagic response targeted to Mm clusters (Fig4 B and C), the fixed 2 dpi infected larvae were imaged by confocal microscopy with a 40X water immersion objective (NA 0.8) over the whole caudal hematopoietic tissue (CHT) region. The same approach was used to quantify Mm acidification in the CHT region (Fig. 6 A, B). To investigate the intramacrophage or extracellular localization of bacteria, fixed 2 dpi larvae were again imaged over the CHT as described above, after which the total number of Mm clusters and the number of clusters inside macrophages was counted. To assay cell death, images from fixed 2 dpi larvae were acquired as above, and the number of cells positive for TUNEL staining in the CHT region was counted manually.

### mRNA preparation and injection

*dram1* or *dram1*^Δ19N^ (negative control) RNA was isolated from wild type or *dram1*^Δ19n/Δ19n^ embryos using QIAzol lysis reagent (79306, QIAGEN) and purified with the RNeasy MinElute Cleanup kit (74204, QIAGEN). cDNA synthesis was performed using the iScript cDNA synthesis kit (1708891, BIO-RAD). Full-length *dram1* cDNA and *dram1*^Δ19N^ cDNA was obtained by PCR amplification using Phusion High-Fidelity DNA Polymerase (M0530S, New England Biolabs). The following primers were used: Forward: CTG CGG CGA GAT GTT TTG GTT; Reverse: CAA AAA CAG TGG GAC ATA CAG TGA A. *dram1* or *dram1*^Δ19N^ PCR products were ligated into a ZERO BLUNT TOPO vector (450245, ThermoFisher) and the insert was confirmed by Sanger sequencing (Base Clear, Netherlands). *dram1* and *dram1*^Δ19N^ mRNA was generated using the SP6 mMessage mMachine kit (AM1340, Thermo Fisher) and Poly(A) Tailing Kit (AM1350, ThermoFisher); purified using the RNeasy Min Elute Cleanup kit (74204, QIAGEN) and 50pg mRNA was microinjected into one cell stage embryos.

### TUNEL assay

Cell death was examined by Terminal deoxynucleotidyl transferase dUTP nick end labelling (TUNEL staining) with the In Situ Cell Death Detection Kit, TMR red (1256792910, SIGMA-ALDRICH) in 2dpi fixed embryos. The assay was performed as follows: embryos were fixed in 4%PFA solution O/N at 4 °C, de-hydrated and re-hydrated in serial dilutions of methanol (MeOH) (25%, 50%, 75%, 100%, 75%, 50%, 25%) and washed in PBS-TX. Then, embryos were permeabilized in 10 μg/mL Proteinase K for 40 min at 37 °C followed by a quick rinse in PBST. TUNEL staining was performed according kit instructions O/N at 37°C in the dark. Samples were washed and stored in PBST until imaging with confocal microscopy as described above.

### LysoTracker staining and Myeloperoxidase (Mpx) activity assay

Infected embryos were immersed in egg water with 10 μM LysoTracker Red DND-99 (L7528, ThermoFisher) for 1 h. Embryos were washed 3 times with egg water before imaging. Myeloperoxidase (Mpx) activity assay was performed with the Leukocyte detection Kit (390A, SIGMA-ALDRICH) for detection of neutrophils as previously described (73).

### Drug treatment

Embryos were bath treated with Bafilomycin A1 (BafA1) (B1793-10UG, SIGMA-ALDRICH) diluted into egg water at the working concentration of 100 nM for 12h.

### Caspase activity assay

Inflammatory caspase activity was assayed with the fluorometric substrate Z-YVAD 7-Amido-4-trifluoromethylcoumarin (Z-YVAD-AFC, Caspase-1 Substrate IV, Colorimetric, sc-311283, Santa Cruz) as described previously (74). 35 embryos/group were sonicated and incubated in hypotonic cell lysis buffer (*25* mM 4-(2-hydroxyethyl) piperazine-1-ethanesulfonic acid, 5 mM ethylene glycol-bis (2-aminoethyl ether)-N,N,N’,N’-tetraacetic acid, 5 mM dithiothreitol, pH 7.5 on ice for 15 min. For each reaction, 10 μg protein was incubated for 90 min at 28°C with 50 μM YVAD-AFC in 50 μl of reaction buffer (0.2% 3-[(3-cholamidopropyl) dimethylammonio]-1-propanesulfonate (CHAPS), 0.2 M 4-(2-hydroxyethyl) piperazine-1-ethanesulfonic acid, 20% sucrose, 29 mM dithiothreitol, pH 7.5). After the incubation, fluorescence was measured in a Tecan M1000 microplate reader at an excitation wavelength of 400 and emission wavelength of 505 nm.

### Morpholino Injection condition

Previously validated *caspa* and *gsdmeb* morpholinos (MO) (39, 75) were purchased from Gene tools (Gene Tools, USA). MO oligonucleotide sequence: *caspa* 5’-GCCAT GTTTAGCTCAGGGCGCTGAC-3’, *gsdmeb* MO 5’-TCATGCTCATGCTAGTCAGGGAGG-3’ (39, 75). MOs were diluted in Milli-Q water with 0.05% phenol red and 1nL of 0.6 mM (*caspa* MO or 0.7 mM (*gsdmeb*) MO was microinjected into the yolk of one cell stage embryos as previously described (39.)

### Statistical analyses

Statistical analyses were performed using GraphPad Prism software (Version 5.01; GraphPad). All experimental data (mean ± SEM) was analyzed using unpaired, two-tailed t-tests for comparisons between two groups and one-way ANOVA with Tukey’s multiple comparison methods as a posthoc test for comparisons between more than two groups. (ns, no significant difference; *p < 0.05; **p < 0.01; ***p < 0.001). For segregation from F1 or F3 heterozygous, data were analysed with a Chi-square test (ns, no significant difference).

## Acknowledgements

We thank Daniel Klionsky for sharing of the *CMV:EGFP-map1lc3b* transgenic zebrafish line and Georges Lutfalla for the *mpeg1:mCherryF* line. We are grateful to all members of the fish facility team for zebrafish caretaking. We would like to thank Gerda Lamers and Joost Willemse for advice on confocal imaging and image analysis. R.Z. was supported by a grant from the China Scholarship Council (CSC). M.V. and G.F.-C. were funded by European Marie Curie fellowships (H2020-MSCA-IF-2014-655424 and H2020-COFUND-2015-FP-707404), V.T. was a Marie Curie fellow in the Initial Training Network FishForPharma (PITN-GA-2011-289209), and M.v.d.V. was supported by the Netherlands Technology Foundation TTW (project 13259).

## Supplementary figures

**Figure S1:**
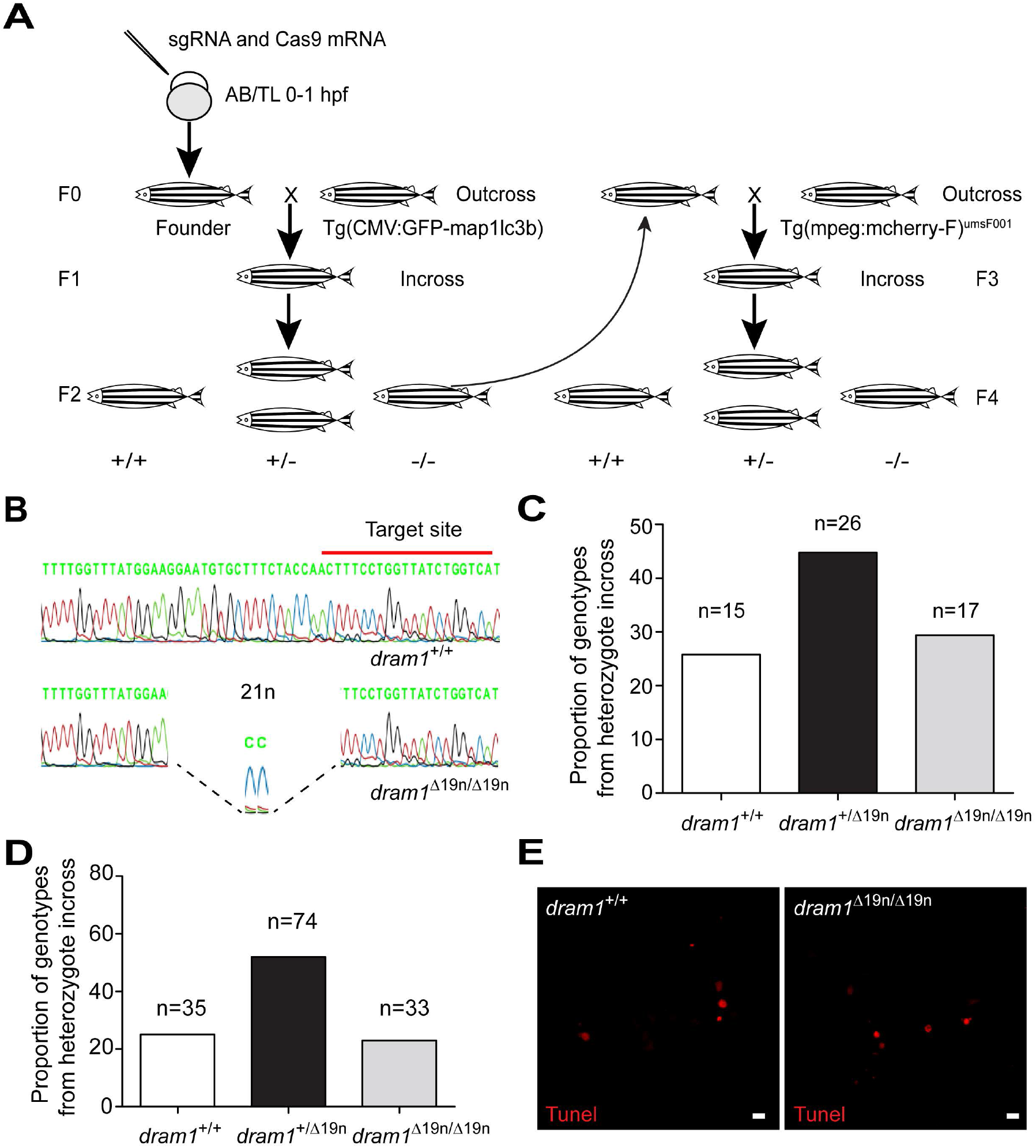
Generation and characterization of the *dram1* mutant line. A. Schematic diagram showing the workflow used for the generation of *dram1* mutant lines. Target-specific sgRNA and Cas9 mRNA were co-injected into one cell stage embryos (AB/TL, wild type line). Founders were outcrossed to *Tg*(*CMV:EGFP-map1lc3b*) or wild type fish to obtain F1. After 3-4 months, the F1 was incrossed to obtain homozygous mutant and wild type F2 siblings. *dram1*^Δ19n/Δ19n^ were outcrossed with the macrophage marker *Tg(mpeg1:mCherryF)^umsF001^* and after 3-4 months subsequently incrossed to obtain *dram1*^+/+^, *dram1^Δ19/+^*, and *dram1*^Δ19n/Δ19n^ carrying *Tg(mpeg1:mCherryF)^umsF001^*. B. Sanger sequencing of dram1Δ19n/Δ19n and dram1+/+ from *F2* offspring. Red lines indicate CRISPR/Cas9 target sites. The genomic DNA was isolated from fin tissue (>3 months old fish). The dram1Δ19n/Δ19n mutant allele has 21 nucleotides deleted and 2 nucleotides inserted. C. Segregation from *dram1*^+/Δ19n^ F1 heterozygous incross. Genotypes of adult fish (>3 months old) were combined from at least three independent breedings and confirmed by PCR and Sanger sequencing. Data were analyzed by Chi Square test. ns, non-significant,*p<0.05,**p<0.01,***p<0.001. D. Segregation from *dram1^Δ19n+^/mpeg1:mCherryF* F1 heterozygous incross. Genotypes of adult fish (>3 months old) combined from at least three independent breedings were confirmed by PCR and sequencing. Data were analyzed by Chi square test. ns, non-significant,*p<0.05,**p<0.01,***p<0.001 E. Representative confocal micrographs of sections from the tail region showing TUNEL staining performed on *dram1*^Δ19n/Δ19n^ and *dram1*^+/+^ larvae at 3dpf. Scale bar, 10 μm.

**Figure S2:**
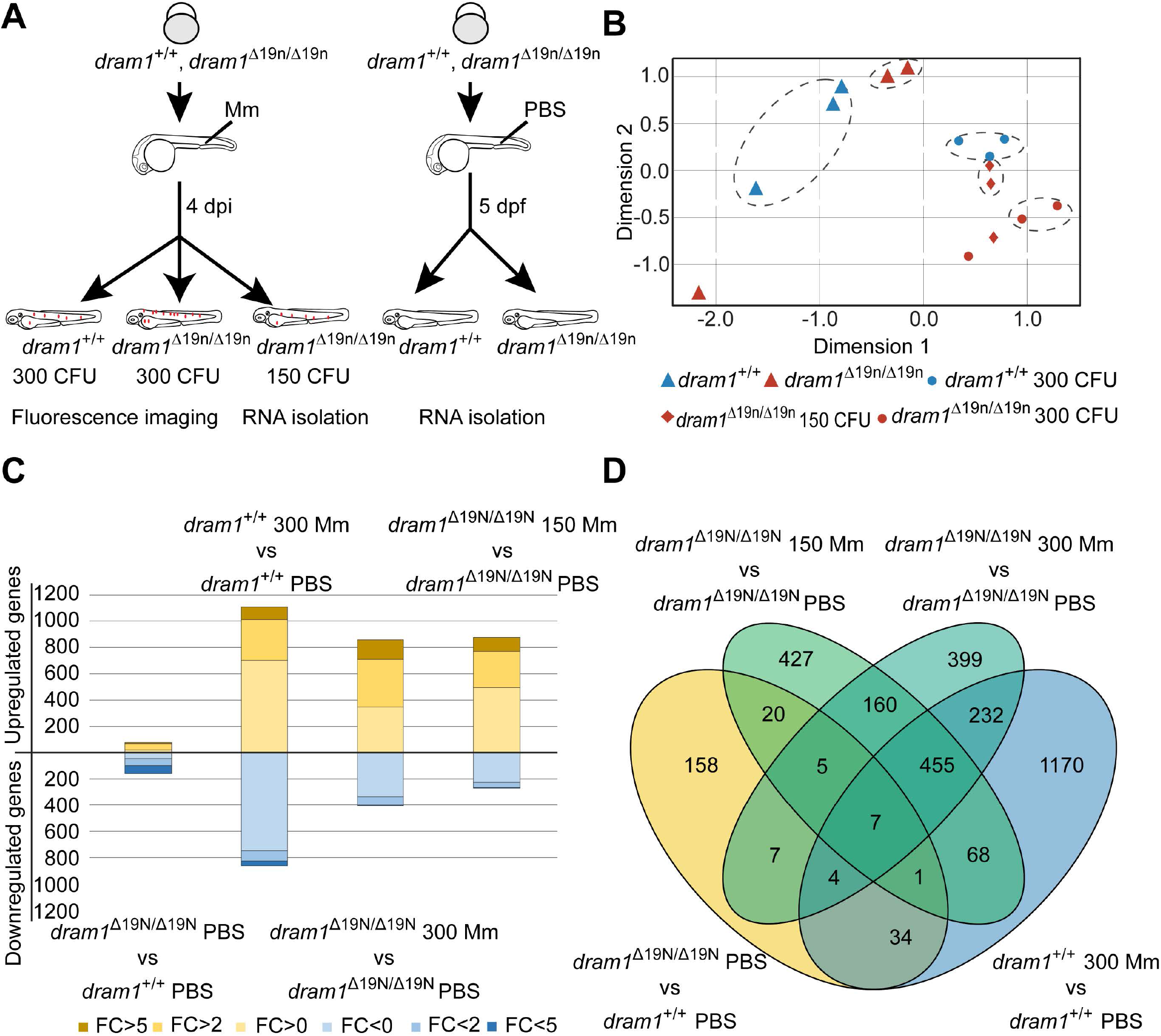
Transcriptome analysis of uninfected and infected *dram1* mutants A. Experimental design to obtain samples for RNA deep sequencing. *Mycobacterium marinum* strain M (Mm) fluorescently labeled with mCherry was microinjected into the blood island of embryos at 28 hpf at an injection dosage of 300 CFU of 150 CFU. Control groups were injected with PBS. B. Principal component analysis of the gene expression data obtained by RNA sequencing. The RNA sequencing samples clustered according to their condition, as pictured by the dashed ellipses grouping the samples. The data sets of one family of *dram1*^Δ19n/Δ19n^ (Mm infected and uninfected) diverged from the rest of the libraries (data points outside the dashed ellipses) and were discarded from the analysis. C. General profile of differential gene expression between the different conditions. The number of genes upregulated are coloured in yellow while downregulated in blue, with indication of the fold-change by colour intensity. D. Venn diagram of the differentially expressed genes common and different between the *dram1*^Δ19n/Δ19n^ and *dram1*^+/+^,dram1^Δ19n/Δ19n^150 CFU and *dram1*^Δ19n/Δ19n^ PBS, *dram1*^Δ19n/Δ19n^ 300 CFU and *dram1*^Δ19n/Δ19n^, *dram1*^+/+^ 300 CFU and *dram1*^+/+^ PBS comparisons.

**Figure S3:**
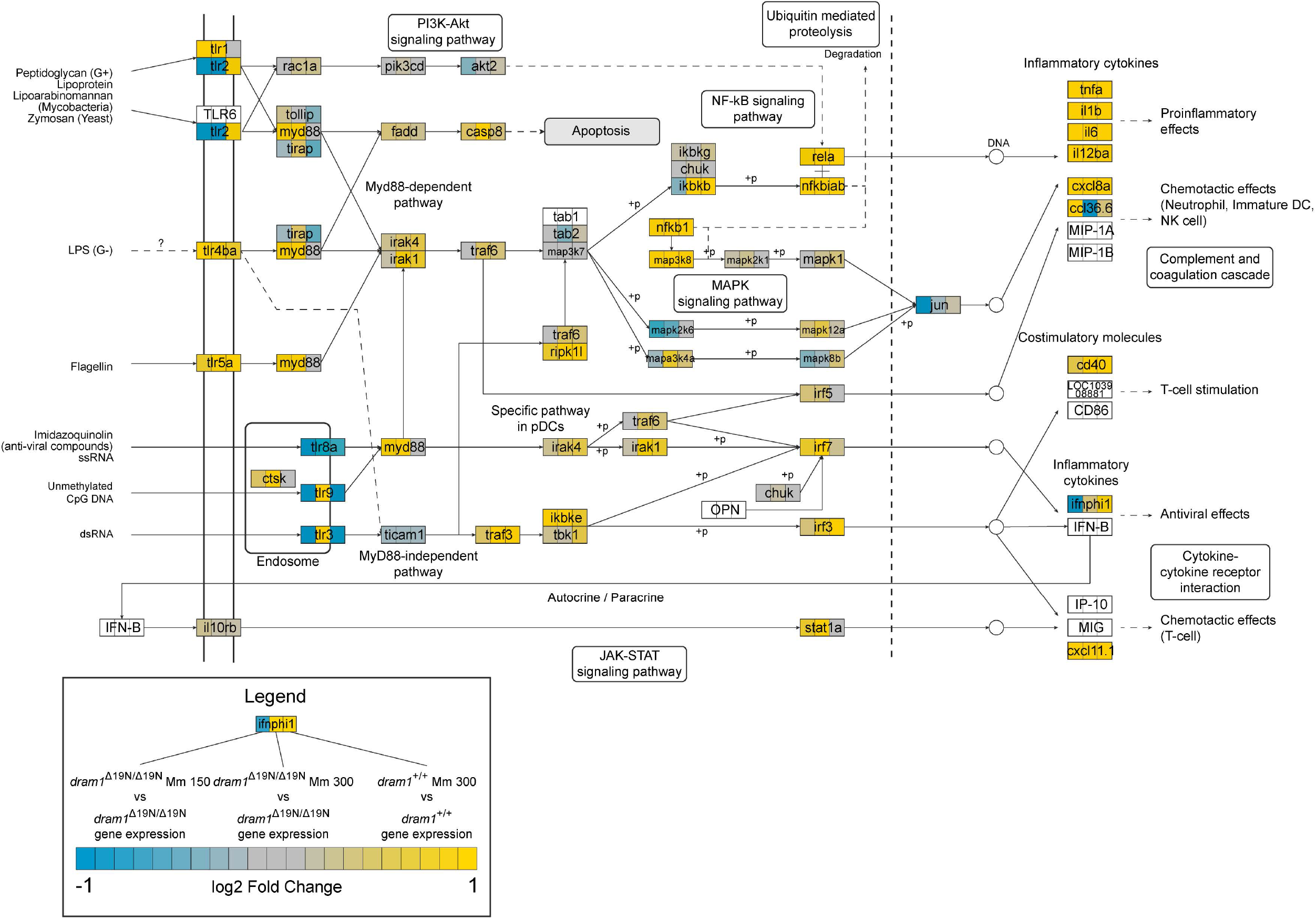
Effect of *dram1* mutation on TLR signaling KEGG pathway of TLR signaling showing differential gene expression in infected *dram1*^Δ19n/Δ19n^ and *dram1*^+/+^. The three data sets used for comparison are shown in the legend of the figure. The expression fold change of the genes is depicted by colour (yellow, upregulated, blue downregulated).

## Supplementary tables

Supplementary Table 1. Enrichment of gene sets altered in *dram1*^Δ19n/Δ19n^ larvae under basal conditions.

A. Gene Ontology categories significantly over and underrepresented in the significant genes differentially regulated between *dram1*^Δ19n/Δ19n^ PBS-injected mutants compared to *dram1*^+/+^ larvae.

B. Gene sets from the MSigDB C2 database significantly positively correlated to the *dram1*^Δ19n/Δ19n^ mutants transcriptome compared to *dram1*^+/+^ larvae.

C. Gene sets from the MSigDB C2 database significantly negatively correlated to the *dram1*^Δ19n/Δ19n^ mutants transcriptome compared to *dram1*^+/+^ larvae.

For data set see: https://doi.org/10.5281/zenodo.2615900

**Supplementary Table 2.** Zebrafish lines used in this study

| Name | Description | Reference |
| --- | --- | --- |
| AB/TL | Wild type strain | 6 |
| *Tg(CMV:EGFP-map1lc3b)* | GFP reporter transgenic zebrafish for Lc3 | 26 |
| *Tg(mpeg1:mCherryF)<sup>umsF001</sup>* | Macrophage marker with membrane-localizing *mCherryF* | 27 |
| *dram1<sup>+/+</sup>/GFP-Lc3* | Siblings of *dram1<sup>ibl53</sup>* carrying a transgenic GFP-Lc3 reporter | In this study |
| *dram1<sup>Δ19n/Δ19n</sup>/GFP-Lc3* | *dram1<sup>ibl53</sup>* mutant line (Δ19n indel) carrying a transgenic GFP-Lc3 reporter | In this study |
| *dram1<sup>+/+</sup>/mpeg1:mCherryF* | Siblings of *dram1<sup>ibl53</sup>* mutant line carrying a transgenic *mpeg1:mCherryF* reporter | In this study |
| *dram1<sup>Δ19n/Δ19n</sup>/mpeg1:mCherryF* | *dram1<sup>ibl53</sup>* mutant line (Δ19n indel) carrying a transgenic *mpeg1:mCherryF* reporter | In this study |

**Supplementary Table 3.**
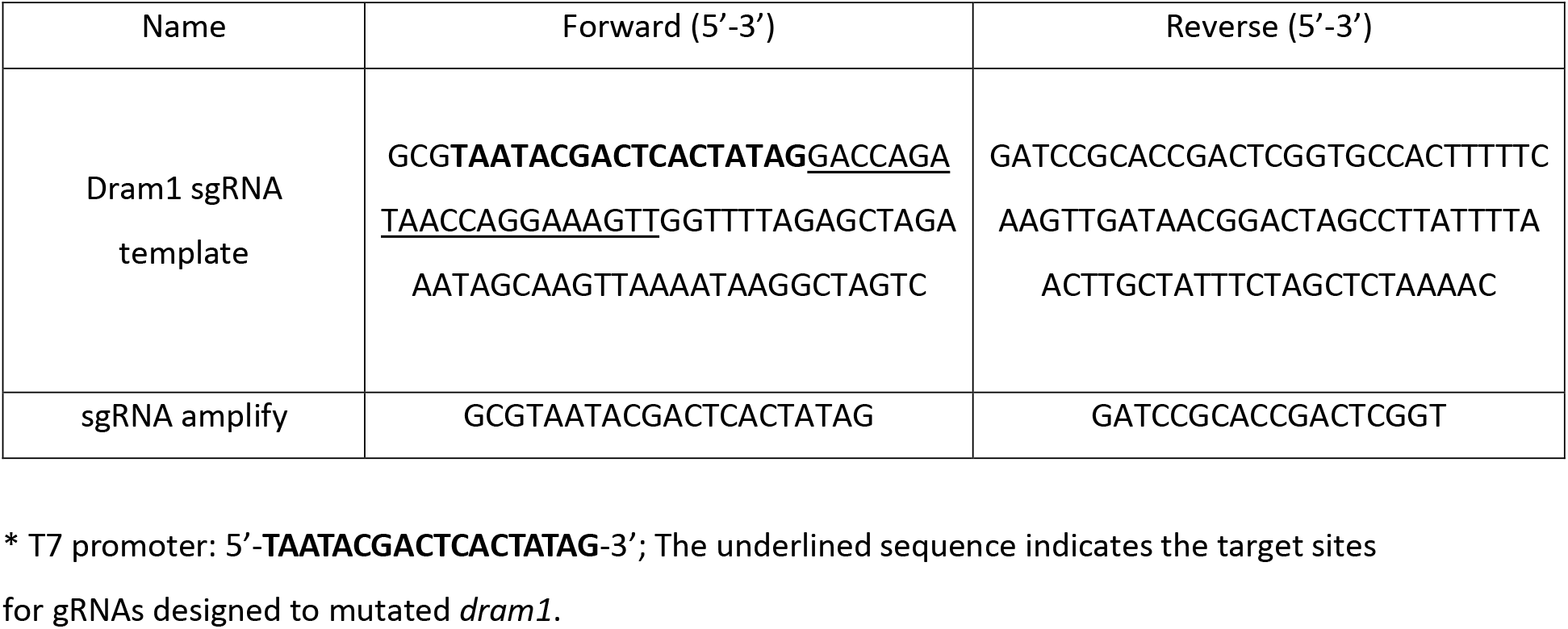
Primers for complementation and amplification of sgRNA

